# AmpliPiper: A versatile amplicon-seq analysis tool for multilocus DNA barcoding

**DOI:** 10.1101/2024.12.11.628038

**Authors:** Astra Bertelli, Sonja Steindl, Sandra Kirchner, Paula Schwahofer, Elisabeth Haring, Nikolaus Szucsich, Luise Kruckenhauser, Martin Kapun

## Abstract

The advent of third generation sequencing technology has revolutionized parallelized sequencing of DNA fragments of varying lengths, such as PCR amplicons, which provides unprecedented new opportunities for large-scale and diverse DNA barcoding projects that, for example, aim to quantify the accelerating biodiversity crisis. However, the broad-scale application of these new technologies for biodiversity research is often hindered by the demand for advanced bioinformatics skills to carry out quantitative analyses. To facilitate the application of multilocus amplicon sequencing (amplicon-seq) data for biodiversity and integrative taxonomic research questions, we present AmpliPiper, an automated and user-friendly software pipeline which carries out bioinformatics analyses of multilocus amplicon-seq data generated with Oxford Nanopore (ONT) sequencing. AmpliPiper combines analysis methods for DNA barcoding data that include demultiplexing of pooled amplicon-seq data, haplotype-specific consensus sequence reconstruction, species identification based on comparison to the BOLD and GenBank databases, phylogenetic analyses and species delimitation. We demonstrate the applicability and workflow of our approach based on a newly generated dataset of 14 hoverfly (Syrphidae) samples that were amplified and sequenced at four marker genes. We further benchmark our approach with Sanger sequencing and simulated amplicon-seq data which show that DNA barcoding with ONT is both accurate and sensitive to detect even subtle genetic variation.

## 1 Introduction

The quantitative characterization of organismal diversity is of central importance for depicting and understanding the extent and ultimate causal reasons for biodiversity loss in the face of an accelerating global biodiversity crisis and the dramatic habitat loss due to extensive agriculture, man-made pollution and ongoing climate change (Harfoot et al., 2021; Hopkins & Freckleton, 2002; IPBES, 2019; Isbell et al., 2023). Recent advances in DNA sequencing technology facilitate the application of molecular genetic methods in support of a purely morphological characterization of species identity and taxon diversity (Gajski et al., 2024; Scarano et al., 2024). DNA barcoding, where taxa are identified based on their unique DNA sequence, ideally at a single marker locus (Hebert et al., 2003), has gained enormous diagnostic power due to the availability of ever-growing reference databases such as the Barcode of Life Database (BOLD) by the international iBol consortium (Ratnasingham et al., 2024).

Haploid standard markers, such as the mitochondrial (mt) cytochrome c oxidase subunit 1 gene (COI or COX1) in animals or the 12S (12S) and the 16S rRNA genes (16S), the latter in particular used in bacteria, are well-established (for reviews, see Antil et al., 2023; Chase & Fay, 2009; Gostel & Kress, 2022; Kress & Erickson, 2012; von Cräutlein et al., 2011). However, while well applicable in most groups, this approach is often insufficient in taxon groups with complicated systematics (discussed, e.g., in Hawlitschek et al., 2017; Hebert et al., 2003; Liu et al., 2017). Different sources of error can hamper identifications solely based on single haploid markers. For example, since mitochondria are inherited maternally in most species, mitochondrial marker genes do not allow to detect hybridizations (e.g., Dejaco et al., 2016; Ermakov et al., 2015; Zakharov et al., 2009). The same applies to single haploid markers of plastomes in plants, where the uniparental inheritance prohibits tracking hybridization events, which often play a pivotal evolutionary role (Fazekas et al., 2012; Hollingsworth et al., 2016). Single marker approaches are likewise prone for error in cases of recent speciation events, when incomplete lineage sorting can disguise actual species borders (Degnan & Rosenberg, 2009; Maddison, 1997; Nabholz, 2024). In general, DNA barcoding has its limitation whenever intraspecific variation is not significantly lower than interspecific variation, a circumstance which might be gene-specific due to differences in the evolutionary rate among genes (e.g., Krawczyk et al., 2014). Another common problem of mt markers is the possible occurrence of NUMTs, i.e., copies of full or partial mt genes (or sometimes even complete mt genomes) integrated in the nuclear genome (Hebert et al., 2003). Such paralogs are on the long run characterised by pseudogenization and their evolutionary rates differ from the rates of their ancestor genes in the organelle genomes (Perna & Kocher, 1996). Thus, NUMTs can impair both correct species identification through DNA barcoding as well as phylogenetic inference and can ultimately bias estimates of species richness (Hebert et al., 2023; Song et al., 2008). Besides NUMTs other paralogous sequences, such as duplicated genes and pseudogenes, but also chimeric sequences (e.g., PCR artefacts), polyploidy, contamination, and reduced amplification success due to partially degraded DNA affect success and reliability of DNA barcoding, even more, when only single loci are used (e.g., Ermakov et al., 2015; Krawczyk et al., 2014). For these reasons, multi-locus barcoding approaches, which became more popular recently, gain diagnostic power by integrating evolutionary signals across various markers and have proven more robust for both species identification and delimitation (e.g., Klimov et al., 2019; Mallo & Posada, 2016). However, the amplification and sequencing of multiple markers with classical Sanger sequencing, which is still considered the gold standard for DNA barcoding, is often limited because of cost and time constraints (Shokralla et al., 2014). Moreover, sequencing nuclear markers of heterozygous organisms often necessitates additional cloning steps for accurate separation of different haplotypes (Kress & Erickson, 2012).

Some of the limitations in parallelization can be overcome by next generation sequencing technologies, which allow to generate myriads of DNA sequences in one sequencing run. Illumina sequencing, which yields highly accurate reads, is currently limited by read lengths and cannot produce sequences longer than 250 bp. This complicates the reconstruction of phased haplotypes of amplicons longer than the lengths of partially overlapping read pairs. However, this limitation is alleviated by the recent advent of third generation sequencing technologies such as Oxford Nanopore sequencing (ONT) or Pacific Bioscience sequencing (Gajski et al., 2024). These technologies now allow to sequence long DNA fragments of hundreds of kilo-base pairs length in a highly parallelized fashion (Scarano et al., 2024). They thus help to overcome several major obstacles that previously hindered a parallelized application of DNA barcoding to large samplesets and allow for combined analyses of multiple fragments. Particularly ONT sequencing has gained popularity and is nowadays commonly used for small to medium-sized sequencing projects (Baloğlu et al., 2021; Hebert et al., n.d.; Koblmüller et al., 2024; Srivathsan et al., 2018) due to fairly simple library preparation protocols, the small size of the MinION sequencing device, which can be operated on any benchtop computer (Lu et al., 2016), and the comparably small sequencing costs (Satam et al., 2023). However, in spite of these advantages, ONT sequencing is still characterised by a high error rate of up to 5-10%, even with the most recent library preparation chemistry and pore types (R10.4; Ni et al., 2023). Thus, advanced and sophisticated bioinformatic tools are needed to account for sequencing errors in these datasets.

This resulted in a recent emergence of various software solutions for processing amplicon sequencing data, which address different steps of the bioinformatic analysis pipeline underlying DNA barcoding with NGS data. ONTBarcoder (Srivathsan et al., 2018), for example, is a bioinformatics tool specifically designed for haploid single-locus markers based on ONT sequencing data. It is able to process both already base-called, but also raw real-time sequencing data, and reconstructs the barcodes through iterative consensus building. NGSpeciesID (Sahlin et al., 2021), which implements several software tools and allows clustering, consensus sequence reconstruction, and polishing of amplicon sequencing data of single loci, but requires that the data are filtered (for size and/or quality) and demultiplexed by locus, prior to running the program. EMU (Curry et al., 2022), is specifically tailored for microbial 16S sequences from long-read metagenomic samples, and limited to single-locus approaches. EMU also offers the possibility to perform taxonomic identification from a custom database. Decona (Oosterbroek et al., 2021), likewise combining several tools, was specifically developed to tackle the challenges of eDNA/bulk sequencing, and also allows processing multi-locus data. While all of these software solutions have proven powerful for the analyses of amplicon sequencing data, they are limited because they generally do not account for different haplotypes in di- and polyploid organisms, when using nuclear instead of haploid mitochondrial or plastid markers.

Conversely, amplicon_sorter (Vierstraete & Braeckman, 2022), a standalone python script specifically tailored to ONT amplicon sequencing data, performs alignment-based consensus sequence reconstruction leveraging fast and iterative clustering of reads and is sensitive enough to distinguish different haplotypes in multi-locus input data even without prior demultiplexing.

Taxonomic identification, as one of the main tasks in DNA barcoding, is however not included in all of the aforementioned software tools and thus requires subsequent manual comparison of the consensus sequences to databases such as BOLD or NCBI GenBank. Only EMU offers an automated identification step, however, the software requires constructing a local reference database a priori. Moreover, additional phylogenetic analyses, despite being crucial to support and inform taxonomic research based on morphological features, are not included in the aforementioned software tools and hamper an integrative taxonomic approach. Especially the advanced bioinformatic skills, required from users to combine different software tools may hinder a broader application of ONT sequencing and a full exploitation of the extensive potential of this technology for biodiversity research.

To facilitate and simplify bioinformatic analyses of DNA barcoding data, in particular of multiplexed multilocus PCR amplicons, we developed AmpliPiper, a fully automated open-source analysis pipeline written in the BASH scripting language, Python and R, which is available at https://github.com/nhmvienna/AmpliPiper. Our approach combines both open-source third party software and customly written scripts to conduct highly automated multilocus analyses of multiplexed PCR amplicons that were generated with ONT sequencing. AmpliPiper uniquely integrates read filtering and locus-specific demultiplexing based on information from primer sequences of the multiplexed loci. Locus- and sample-specific consensus sequence reconstruction are then carried out for different haplotypes using amplicon_sorter (Vierstraete & Braeckman, 2022) described above. Based on these results, AmpliPiper estimates the ploidy level at each locus using a maximum likelihood approach. Subsequently, AmpliPiper combines all consensus sequences across samples for each locus, generates multiple alignments and carries out phylogenetic analyses. In addition, AmpliPiper employs comparisons of consensus sequences from common barcoding loci to the BOLD or NCBI GenBank databases for species identification and performs locus-wise species delimitation analyses using the ASAP (Puillandre et al., 2021) approach. To facilitate debugging, AmpliPiper generates verbose log files and comprehensive reports and displays all final analysis results summarised in a HTML file. All output data generated from the different analyses steps are provided in a well-structured output folder and can be used for subsequent analyses. We demonstrate the workflow and performance of AmpliPiper using both simulated ONT data and a newly generated dataset of ONT amplicon sequencing data of four maker genes from 14 individuals of hoverflies (Syrphidae) from three different subtribes and discuss the workflow and performance of our approach, but also unresolved challenges and current limitations of multi-locus AmpliSeq with ONT sequencing technology.

## 2 Methods

### 2.1 Installation

#### 2.1.1 Native installation

AmpliPiper with native installation has been successfully tested on various Linux distributions, including AlmaLinux (v.8) and Ubuntu (v22.04). The software package managers *conda* and *mamba*, as well as *git* utilities, need to be preinstalled to download and locally install all third-party software required for the AmpliPiper pipeline. The installation of the required programs can be started manually by executing the *setup.sh* script in the shell folder or will be automatically executed the first time AmpliPiper is started. All necessary third party programs are locally installed within the AmpliPiper folder as separate *conda* environments, which are sequentially loaded during the AmpliPiper analysis pipeline.

#### 2.1.2 Docker installation

To enhance the portability of AmpliPiper across various operating systems, we developed a Docker image containing all necessary dependencies, allowing AmpliPiper to run consistently across different platforms. This Docker image is built upon *condaforge/miniforge3*, which inherently includes *conda* and *mamba* executables and complies with the installation prerequisites. We loaded the pipeline scripts inside the image according to the local file system’s structure and executed the *setup.sh* script to recreate the *conda* environments within the Docker image. The image building and pushing process is automated through a continuous integration/continuous delivery (CI/CD) workflow configured with GitHub Actions. This setup ensures that a new image is built and updated with every code change committed to the repository. The Docker image is available under the GitHub Container Registry and can be accessed via the URL: <u>ghcr.io/nhmvienna/amplipiper</u>. We have verified the installation workflow on Windows (version 10.0.22631.4460) and Ubuntu (version 22.04), and it should be compatible with all AMD64-based platforms.

### 2.2 Input files

The AmpliPiper workflow requires two comma-separated files as primary input: (1) The first input file (PRIMERS) provides metainformation on the multiplexed amplified loci that should be demultiplexed in the sequencing dataset. The data in each row represents a different locus and has to include the name of the locus, the locus-specific forward and reverse primer sequences and the locus-specific expected fragment-size. (2) The second file (SAMPLES) provides the sample names and the file paths to the raw FASTQ files of all samples in the dataset. These raw FASTQ files have to be demultiplexed per sample, e.g. based on sequencing barcodes, prior to the AmpliPiper analysis. AmpliPiper will subsequently further demultiplex the FASTQ for the different loci in the PRIMERS file per specimen.

### 2.3 AmpliPiper workflow

#### 2.3.1 Primer similarity

Before starting the demultiplexing and consensus reconstruction, AmpliPiper evaluates the genetic distance among all possible primer sequences in the PRIMERS input and their reverse complements with a custom script (*CompPrimers.py*) that utilises the *edlib* Python package (Šošić & Šikić, 2017). Primer sequences of different loci which are highly similar may confound the demultiplexing of raw FASTQ files according to loci as part of the AmpliPiper pipeline. The output from the analysis of primer sequence similarity can thus be used to adjust the sensitivity of the subsequent demultiplexing step which clusters reads based on similarity to primer sequences (see below).

#### 2.3.2 Filtering and demultiplexing of loci

First, all raw reads are filtered by minimum average base quality with *Chopper* (De Coster & Rademakers, 2023). Then, the raw reads in each sample-specific FASTQ file are demultiplexed based on the information in the PRIMERS input file using a custom Python script (*demultiplex.py*). Based on the *edlib* Python package (Šošić & Šikić, 2017), this script tests for sequence similarity of forward and reverse primer sequences (and their reverse complements) at the beginning and end of each read. Only reads with high sequence similarity with both primers and whose fragment lengths are within the expected size range are assigned to a given locus. As a result, this script returns locus-specific reads for each sample. The maximum number of reads that are subsequently used for consensus sequence reconstruction can be defined by the user. If this threshold is smaller than the total number of available demultiplexed reads, then only a subset of the total reads with the highest average base qualities is used for subsequent analyses.

#### 2.3.3 Consensus sequence reconstruction and multiple alignment

Locus- and sample-specific consensus sequences are reconstructed with *amplicon_sorter* (Vierstraete & Braeckman, 2022) based on reads that were pre-filtered and sorted during the demultiplexing step described above. Specifically, the sensitivity of amplicon_sorter to consider similar clusters of reads separately for consensus sequence reconstruction can be manually adjusted. The parameter --similar-consensus defines the threshold similarity among clusters. If the similarity of two clusters is smaller or equal to this threshold, the two clusters are considered separately, otherwise they are collapsed prior to consensus reconstruction. All consensus sequences identified for each locus and sample are further analyzed with the custom Python script *ParseSummary.py*, which summarizes the log files of *amplicon_sorter* that contain information about the numbers of reads supporting each consensus sequence. In a next step, the script *ChooseCons.py* uses the information from the counts of raw reads that support each consensus sequence to test for the ploidy level with the highest likelihood by fitting observed and expected frequencies of different ploidy levels (diploidy, triploidy and tetraploidy) with multinomial probability mass functions to calculate individual likelihoods for each ploidy level. The ploidy with the highest likelihood will subsequently be reported for each locus and sample. All consensus sequences are subsequently combined across samples for each locus and further used for downstream analyses. In addition, all consensus sequences predicted by *amplicon_sorter* are also stored in the output and allow more in-depth and custom analyses outside the AmpliPiper analysis pipeline. After that, both the number of successfully reconstructed consensus sequences that are used for further analyses and the cases where the consensus sequence reconstruction failed are visualised for all samples and loci as a heatmap in R with the custom script *MissingDataHeatmap.r*. Finally, the software *mafft* (Katoh & Standley, 2013) is used to generate multiple sequence alignments (MSA) for each locus in the dataset and the Python library *pymsaviz* implemented in the script *msaviz.py* produces alignment plots for each locus to visually evaluate the alignment quality.

#### 2.3.4 Species identification tools

The application programming interface (API) of the BOLD database (Ratnasingham et al., 2024) as implemented in the custom Python script *BOLDapi.py* is used for species identification whenever the dataset contains consensus sequences of either mt COX1, Internal Transcribed Spacers (ITS) of the nuclear rRNA gene cluster (ITS1 and ITS2), Maturase K and/or Ribulose-1,5-Bisphosphate Carboxylase (MATK_RBCL). This analysis returns the ten best hits with the highest similarity scores in tabular output for each consensus sequence, whenever available. Subsequently, the sample names in all downstream analyses are automatically supplemented with the best matching species ID and the corresponding sequence similarity in percent. Alternatively, it is possible to use the BLAST API from BioPython (Cock et al., 2009) as implemented in the custom script *BLASTapi.py* for species identification. In this case the consensus sequences of either of the above-mentioned loci are compared against the *core_nt* database of NCBI Genbank using the *blastn* algorithm (Camacho et al., 2009). In addition, NCBI Entrez is used to obtain the genus and species names based on the Genbank ID’s of the 10 best BLAST hits for each sequence. Since NCBI Entrez requires authentication via email address, it is necessary to provide an email address when using the BLAST option (e.g., --blast).

#### 2.3.5 Genetic distance, phylogenetic analysis and species delimitation

First, the custom script *GeneticDist.r*, calculates pairwise genetic distances based on the MSAs using the R package *ape* (Paradis, 2006) among all samples for each locus separately and visualises the distance matrices as heatmaps. Then, IQ-tree2 (v2.3.6, Minh et al., 2020) is used for inferring the best-fitting substitution model for each locus separately and for all concatenated loci prior to calculating maximum likelihood phylogenetic trees with 1,000 rounds of bootstrapping for each locus and for all haplotypes concatenated per sample. For the latter dataset, it is optionally possible to estimate substitution rates for each partition, i.e., locus, separately by setting the flag *--partition*. Complementary to this, wASTRAL as implemented in ASTER (v1.19.4.6, Zhang & Mirarab, 2022) is used to construct consensus trees based on the individual trees reconstructed for each locus. Note that phasing across loci is not possible and haplotypes are thus randomly concatenated across loci for an individual, which may confound the analyses of F1 hybrids in the datasets. In addition, the phylogenetic trees are based on the automated alignments of the consensus sequences generated by *mafft* as described above, which may contain alignment errors. We therefore add a “preview” watermark to each phylogenetic plot, generated with *ggtree* (Yu et al., 2017) in R, which should indicate that these trees are preview-only and need to be critically evaluated prior to biological interpretation. Optionally, it is possible to define one or multiple samples from the samples.csv input file as outgroups using the *--outgroup* parameter. This information is then used for rooting the phylogenetic trees. However, if no outgroup is defined or the outgroup sample is missing in the MSA, midpoint rooting is used for plotting the trees. Finally, the MSAs for each locus and for the concatenated dataset across all loci are used for species delimitation based on a comparison of the genetic distances among haplotypes as implemented in the program *ASAP* (Puillandre et al., 2021), which employs a partition approach based on pairwise genetic distances to identify a “barcode” gap in the distribution of genetic distances. This gap is considered indicative for a speciation event and assumed to be the result of differences between within-species genetic variation mostly driven by coalescent processes and among-species variation driven by an absence of genetic exchange and successive divergence due to disruptive selection and genetic drift.

#### 2.3.6 Output

AmpliPiper generates three types of output: (1) An HTML file (see Figure 2), which is automatically loaded as soon as AmpliPiper has finished the analysis, provides a visual overview of the different analysis steps, including a tabular and visual summary of the Amplicon_Sorter consensus sequence reconstruction, the genetic distances among samples for each loci, a table with putative species names inferred from BOLD or BLAST, the MSAs for each locus, all phylogenetic trees and the clustering of samples based on the species delimitation analysis with *ASAP*. (2) The output folder within the working directory contains the summary of the consensus reconstruction with Amplicon_sorter as a CSV file, the MSAs of all loci as FASTA files, the phylogenetic trees in NEWICK format and plotted as PDFs, the tabular results of the species identification analyses as CSV files and all output files of the ASAP analysis. Finally, (3) the log folder contains detailed log files for all analysis steps of AmpliPiper, which allow for debugging and in-depth evaluation of potential errors and unexpected results. Moreover, all intermediate files, such as the raw output of Amplicon_sorter, are retained and can be used for additional analyses beyond the automated analyses provided by AmpliPiper.

**Figure 1:**
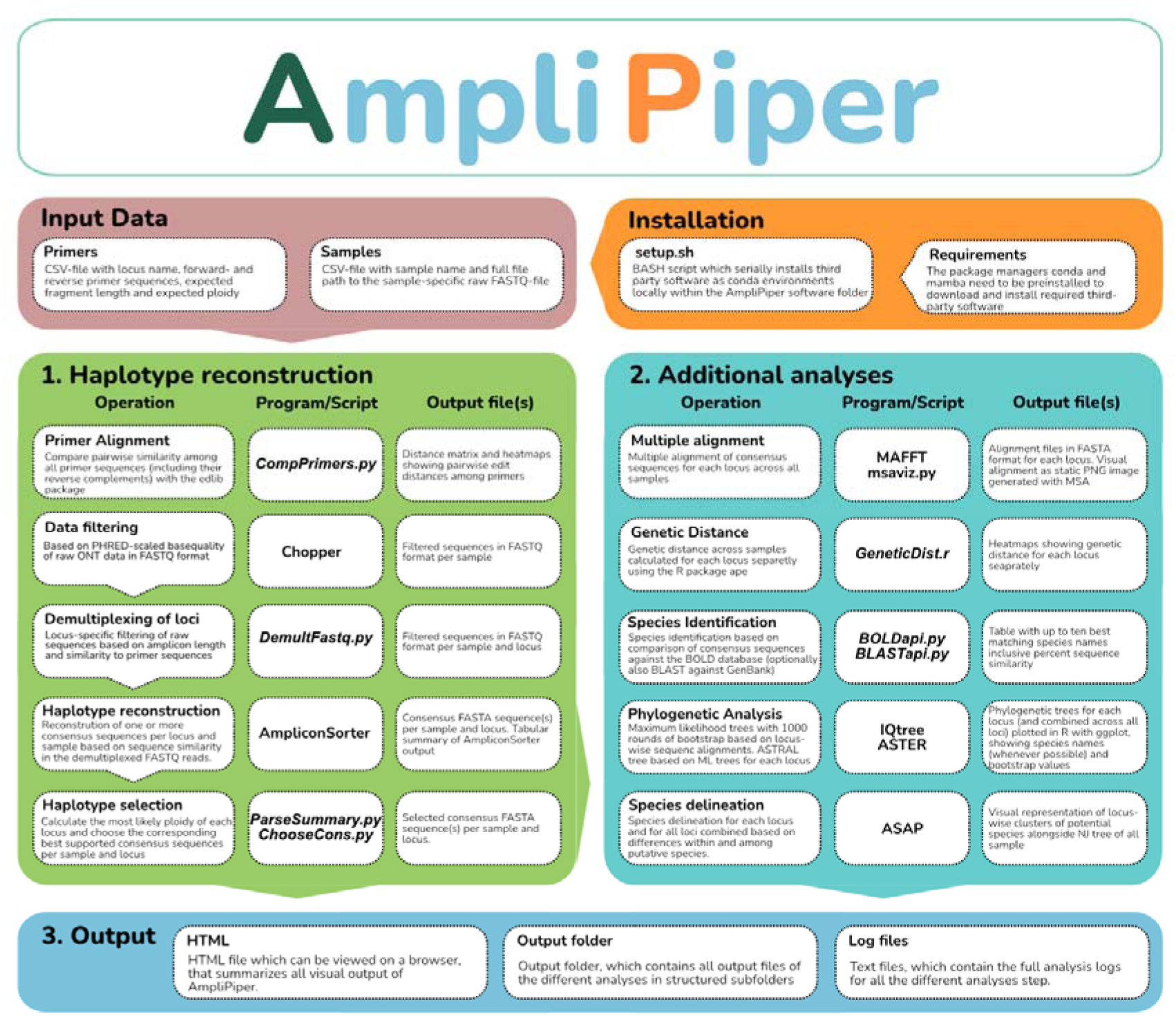
Flowchart depicting the major analysis steps and output files generated by AmpliPiper

**Figure 2.**
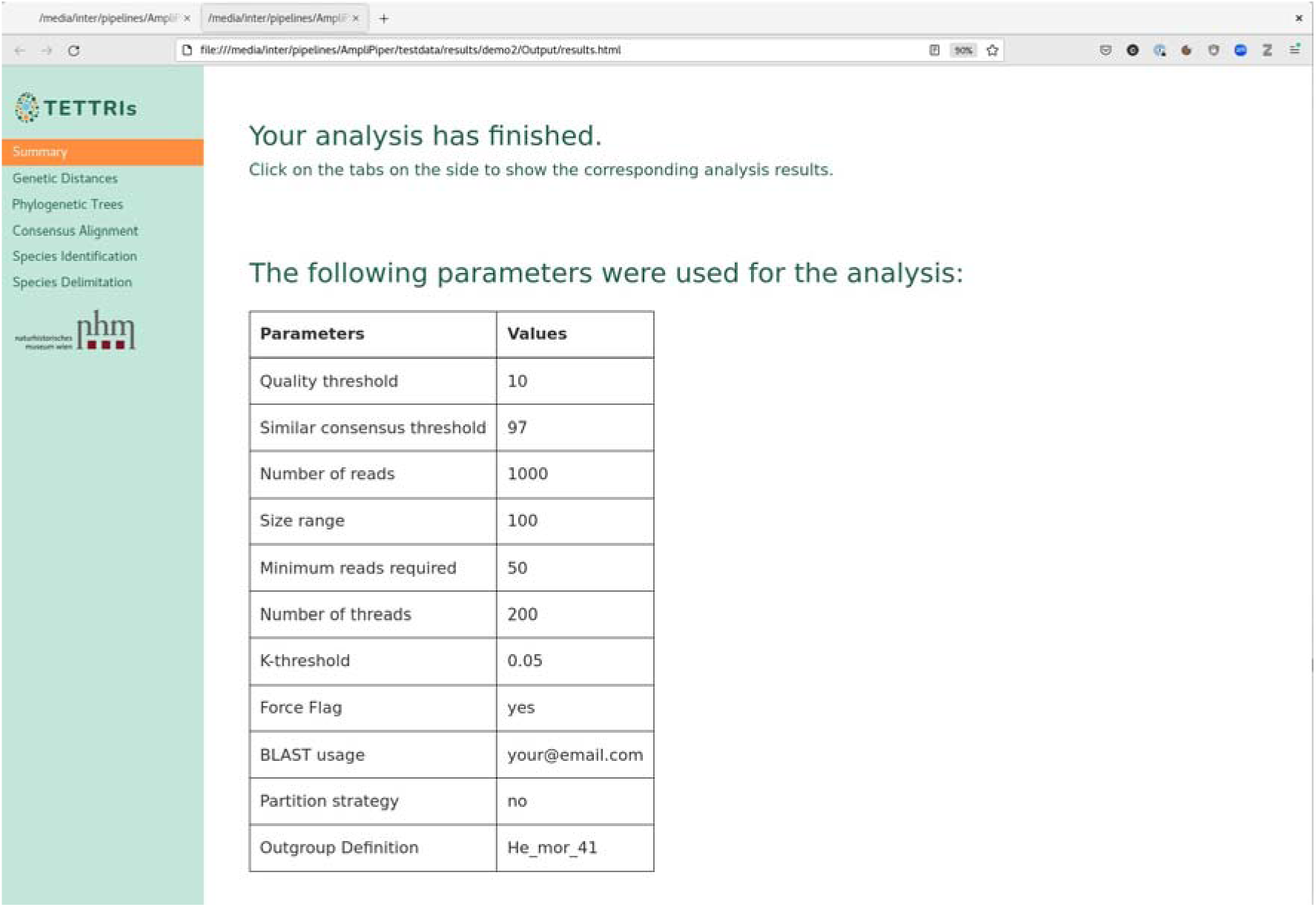
Screenshot of an output HTML page generated by AmpliPiper, which lists the parameters used for the analyses. The sidebar provides links to all the analysis-specific output files.

### 2.4 Test dataset

#### 2.4.1 Material, DNA-extraction, amplification & ONT sequencing

Flies were collected in the course of ABOL BioBlitz actions (Sonnleitner et al., 2022) at different Austrian locations (see Table1). Genomic DNA was isolated following the manufacturer’s protocol of the DNeasy Blood & Tissue Kit (Qiagen). Depending on the size of the specimen, we used one to three legs as input for DNA extraction. For the multilocus amplification, we analysed fragments of four different genes which were employed successfully in a multigene phylogeny of Syrphids (Moran et al., 2022): the mitochondrial **COX1** (mitochondrial cytochrome c oxidase subunit I), **AATS** (Alanyl-tRNA Synthetase 5’ end), **CK1** (Casein Kinase 1 5’ end) and **TULP** (tubby like protein, 5’ end) the same primers were used as described in Moran et al. (2022). In addition, we also amplified a fragment from the nuclear **28S rRNA** gene, the D2-D3 region, using the primer sequences described in Belshaw et al. (2001). We employed a 2-step PCR protocol as described in Pomerantz et al. (2022). In the course of the 1^st^ PCR, amplification was performed in 25 µl reaction volume comprising 12.5 µl Multiplex-Mix from the Multiplex-PCR Kit (Qiagen), 0.5 µM of each primer with entailed ONT universal-tail sequences, 1-2 µl of DNA extract and NfW. The cycling protocol for the 1^st^ PCR was as follows: initial denaturation at 95°C for 10 min, 40 cycles denaturation at 95°C for 45 s, annealing step with ramp from **50°C to 72°** for 1 min 15 s, extension at 72°C for 1 min, and final extension at 72°C for 10 min. We visually checked the results on a 1 % agarose gel and purified the PCR products via paramagnetic beads (AMPure XP, Beckman Coulter Life Sciences) prior to quantifying the concentration with the dsDNA HS Assay Kit for Qbit (ThermoFisher). Based on the resulting DNA concentrations, the PCR products derived from a specimen were pooled in equimolar ratios (10 ng per fragment, 50 ng total) as input for the 2^nd^ PCR.

In the course of the 2^nd^ PCR, the universal tail sequences from the 1^st^ PCR allow the addition of index sequences (individual barcodes) to the end of the amplicons which later on enables the reassignment of sequences to the specimens they originated from. We used the LongAMP Taq 2X Master Mix (NEB) as suggested in the MinION protocol for PCR barcoding of amplicons (SQK-LSK110, ONT) and conducted the 2^nd^ PCR applying the following cycler protocol: initial denaturation at 95°C for 3 min, 15 cycles denaturation at 95°C for 15 s, annealing at 62°C for 15 s, extension at 65°C for 1 min, and final extension at 65°C for 10 min. Again, we checked the PCR products on 1% gel before purification and quantification. The amplicons of all individuals were then pooled in equimolar amounts as input for the library preparation. As suggested by ONT in the Ligation Sequencing protocol for amplicons, we used ∼1 µg (or 100-200 fmol) amplicon DNA as input and followed the manufacturer’s protocol (SQK-LSK114, ONT). The prepared library was then loaded on a Flow Cell (FLO- MIN114) and sequenced for 72 hours on a MinION Mk1C device (MinKNOW version 23.07.12).

After converting the raw ONT sequencing data from POD5 to FAST5 format with the program pod5 (https://github.com/nanoporetech/pod5-file-format), we base-called the reads using guppy (v.6.2.1, Wick et al., 2019) in super accuracy mode based on the “dna_r10.4.1_e8.2_400bps_sup.cfg” model. Subsequently, we further used guppy to demultiplex by sample and trim the raw base-called reads using the EXP-PBC096 barcode sequences. Finally, we visually controlled the quality and length-distribution of the sequencing data with *NanoPlot* (De Coster & Rademakers, 2023).

#### 2.4.1 Sanger sequencing of COX1

To cross-validate our sequences produced with ONT sequencing we generated high-quality reference COX1 barcodes via conventional Sanger Sequencing. Again, amplification was performed in 25 µl reaction volume comprising 12.5 µl Multiplex-Mix from the Multiplex-PCR Kit (Qiagen), 0.5 µM of each primer without the ONT universal-tail (LCO1490, COI-Dipt- 2183R), 1 µl of DNA extract and NfW. The cycling protocol was as follows: initial denaturation at 95°C for 15 min, 45 cycles denaturation at 94°C for 30 s, annealing step at 45°C for 30 s, extension at 72°C for 30 s, and final extension at 72°C for 10 min. The resulting COX1 sequences were sent for sequencing to Microsynth AG and assembled with Geneious Prime 2024.0.3. We then compared these standard DNA barcodes to consensus sequences generated with AmpliPiper. Specifically, we used different thresholds for demultiplexing, ranging from 0 % to 20 % primer sequence differentiation, and a maximum read number used for consensus reconstruction with amplicon_sorter, ranging from 10 to 500 reads, to test for the influence of these parameters on consensus sequence accuracy. We employed the custom Python script *compare2Sanger4sim.py* which uses the *edlib* package for pairwise alignment and comparison and plotted the results in R.

### 2.5 Simulations

To test for the sensitivity and accuracy of our approach in detecting and reconstructing distinct haplotypes within a sample we employed analyses of simulated amplicon-seq data generated with *NanoSIM* (Yang et al., 2017) based on the error profile of our real ONT sequencing data. For the simulations, we used the raw ONT sequencing data of the COX1 locus from sample Sy_tor_30. First we estimated the error profile of our ONT sequencing data for the corresponding sample and locus with the *read_analysis.py* script from the NanoSim package in transcriptome mode. We then artificially introduced random mismatches into the reference sequence using the custom script *mutateRef.py* to obtain synthetic sequences with 1% - 10% sequence differentiation to the original sequence. Based on the results of the error profile obtained before, we then simulated Amplicon-Seq ONT reads with the *simulator.py* script in transcriptome mode using the original Sanger sequence and the artificially mutated sequences as templates and retained aligned simulated reads for further analyses. We subsequently combined random subsets of the simulated sequencing data based on the original and the mutated references at varying frequencies in 20-fold replication and used these data for analysis with AmpliPiper to assess the sensitivity of amplicon_sorter to correctly identify two different haplotypes in diploid datasets with varying levels of differentiation. Specifically, we tested the influence of haplotype frequency (ranging from 10% to 50%) and of read depth (ranging from 100 - 10,000 reads) on the specificity and accuracy of consensus-sequences for each of the two haplotypes that we reconstructed with AmpliPiper. We further inferred the effect of the --similar_consensus parameter of amplicon_sorter (ranging from 96% - 99%), which represents the threshold similarity of independent read clusters that are used for consensus sequence reconstruction. We assessed the specificity of our AmpliPiper approach by counting the number of reconstructed haplotypes for each parameter combination and measured the accuracy, i.e., the number of mismatches between the reference sequences and the reconstructed consensus sequences, using the *edlib* package implemented in the Python script *compare2Sanger4sim.py* and plotted the results in R.

## 3 Results and Discussion

### 3.1 Data analysis of syrphid data with AmpliPiper

Using the amplicon sequencing data from PCR amplifications of one mitochondrial (COX1) and three nuclear markers (CK1, 28S and TULP - AATS was excluded due to too many missing data) for thirteen syrphid specimens and one bombyliid as an outgroup, we employed our AmpliPiper analysis pipeline to reconstruct locus-specific consensus sequences for each individual. Our original MinION sequencing run included additional specimens and loci that were not the primary focus of this study. We therefore constructed a “samples” input file, which consisted of the paths to 14 demultiplexed FASTQ datasets corresponding to the aforementioned focal specimens (Table 1). Furthermore, we included four focal marker loci and their primer sequences in the “primers” file.

**Table 1.** Sampling and barcoding information for the 13 syrphid and one bombyliid specimens used in this study.

| ID | Barcode | IndID | Genus | Species | Date | Location |
| --- | --- | --- | --- | --- | --- | --- |
| Ch_can_39 | barcode51 | Sz22-0103 | *Cheilosia* | *canicularis* | 31.07.2022 | Austria, Carinthia, Malta, meadow near river Malta |
| Ep_bal_38 | barcode50 | Sz22-0120 | *Episyrphus* | *balteatus* | 31.07.2022 | Austria, Carinthia, Malta, meadow near river Malta |
| Er_ten_20 | barcode40 | Sz22-0060 | *Eristalis* | *tenax* | 25.06.2022 | Austria, Lower Austria, Leiser Berge, quarry |
| Er_ten_32 | barcode12 | Sz22-0152 | *Eristalis* | *tenax* | 16.07.2022 | Austria, Carinthia, St. Lorenzen, churchyard |
| Er_rup_34 | barcode14 | Sz22-0229 | *Eristalis* | *rupium* | 09.07.2022 | Austria, Tyrol, Biberwier, lake Weißensee |
| Eu_cor_37 | barcode49 | Sz22-0057 | *Eupeodes* | *corollae* | 24.06.2022 | Austria, Lower Austria, Leiser Berge, Niederleis, Landschaftsteich |
| Eu_cor_22 | barcode42 | Sz22-0062 | *Eupeodes* | *corollae* | 25.06.2022 | Austria, Lower Austria, Leiser Berge, quarry |
| Eu_cor_21 | barcode41 | Sz22-0193 | *Eupeodes* | *corollae* | 08.07.2022 | Austria, Tyrol, NDG Ehrwalder Becken Süd |
| He_mor_41 | barcode53 | Sz22-0247 | *Hemipenthes* | *morio* | 11.06.2022 | Austria, Vienna, Penzing, Paradiesgründe |
| Sc_pyr_35 | barcode15 | Sz22-0063 | *Scaeva* | *pyrastris* | 25.06.2022 | Austria, Lower Austria, Leiser Berge, quarry |
| Sc_sel_50 | barcode62 | Sz22-0150 | *Scaeva* | *selenitica* | 16.07.2022 | Austria, Carinthia, St. Lorenzen, churchyard |
| Sc_sel_49 | barcode61 | Sz22-0222 | *Scaeva* | *selenitica* | 09.07.2022 | Austria, Tyrol, Biberwier, lake Weißensee |
| Sy_tor_30 | barcode10 | Sz22-0073 | *Syrphus* | *torvus* | 25.06.2022 | Austria, Lower Austria, Leiser Berge, Waldweide |
| Sy_tor_31 | barcode11 | Sz22-0151 | *Syrphus* | *torvus* | 16.07.2022 | Austria, Carinthia, St. Lorenzen, churchyard |

In a first analysis step, we compared the minimum pairwise sequence difference among all primer sequences and their reverse complements, since high levels of sequence similarity among certain primers may confound demultiplexing of locus-specific reads. If the kthreshold parameter in our demultiplexing approach is set too liberal, it may not be possible to distinguish between similar primers which may lead to incomplete demultiplexing of the raw reads and subsequently to problems during consensus sequence reconstruction. As shown in Figure SXXX, we found that all primers were highly different and that the 28S_rev and 28S_fwd primers were the most similar pair with a sequence difference of 0.3438, i.e. 34.38% mismatches. Since we used the default kthreshold value of 0.05, which allows no more than 5% mismatches between a given primer and the flanking region of a raw read, we assume that the accuracy of the demultiplexing process is not confounded by primer similarity.

As shown in Table 2, the total number of reads per sample (demultiplexed with guppy v.6.2.1) that passed a minimum average quality threshold of PHRED ≥ 10, ranged from 9,662 (Er_ten_32) to 59,865 (Eu_cor_21). For each of the loci, we further set the maximum number of reads used for amplicon_sorter to 1,000 (--nreads 1000). Of the markers CK1, 28S and COX1, we were able to obtain 1,000 reads unambiguously assigned to the corresponding locus during the locus-specific demultiplexing step either for all individuals (CK1 and 28S) or the large majority of samples (COX1). Yet, for the TULP locus we retrieved 1,000 reads only for 6 out of the 14 individuals . For some specimens, we even obtained less than 100 unambiguous reads for this locus. The PCR for TULP worked well for all specimens, which may suggest that either the index PCR (2^nd^ PCR) amplification of this locus or the subsequent bioinformatic demultiplexing was less efficient compared to the other loci.

**Table 2.** Summary of raw read counts (Total) and demultiplexed locus-specific reads that were used for consensus sequence reconstruction with amplicon_sorter.

| ID | Total | 28S | CK1 | COX1 | TULP |
| --- | --- | --- | --- | --- | --- |
| Ch_can_39 | 28126 | 1000 | 1000 | 1000 | 114 |
| Ep_bal_38 | 33338 | 1000 | 1000 | 468 | 381 |
| Er_rup_34 | 19211 | 1000 | 1000 | 1000 | 64 |
| Er_ten_20 | 28252 | 1000 | 1000 | 1000 | 1000 |
| Er_ten_32 | 9662 | 1000 | 1000 | 419 | 151 |
| Eu_cor_21 | 59865 | 1000 | 1000 | 1000 | 1000 |
| Eu_cor_22 | 59821 | 1000 | 1000 | 1000 | 1000 |
| Eu_cor_37 | 11212 | 1000 | 1000 | 1000 | 636 |
| He_mor_41 | 47278 | 1000 | 1000 | 1000 | 168 |
| Sc_pyr_35 | 14554 | 1000 | 1000 | 1000 | 1000 |
| Sc_sel_49 | 11869 | 1000 | 1000 | 1000 | 900 |
| Sc_sel_50 | 11021 | 1000 | 1000 | 1000 | 814 |
| Sy_tor_30 | 22025 | 1000 | 1000 | 1000 | 1000 |
| Sy_tor_31 | 25086 | 1000 | 1000 | 1000 | 1000 |

#### 3.1.1 Haplotypes and modelled ploidy

Next, we used the demultiplexed reads for consensus sequence reconstruction of each locus and sample. Following the recommendation in the amplicon_sorter manual for ONT data generated with R10.4 pores, we increased the default “--similar-consensus” parameter from 96% to 97%. We obtained a single specific consensus sequence from each sample for the loci 28S, COX1 and CK1 when removing reconstructed haplotypes that are supported by less than 10% of all raw reads, which was expected for the supposedly haploid mitochondrial COX1 locus, but not for the nuclear loci 28s and CK1. These results may either suggest that the individuals in our study were highly inbred and homozygous at these loci or that these two loci are generally not very variable. Low levels of variability, i.e., only few differences between haplotypes, may be too subtle to be identified by amplicon_sorter in the error-prone ONT sequencing data. The latter hypothesis is supported by the consensus reconstruction results for the TULP locus, which yielded three heterozygous individuals (see Figure 3).

**Figure 3.**
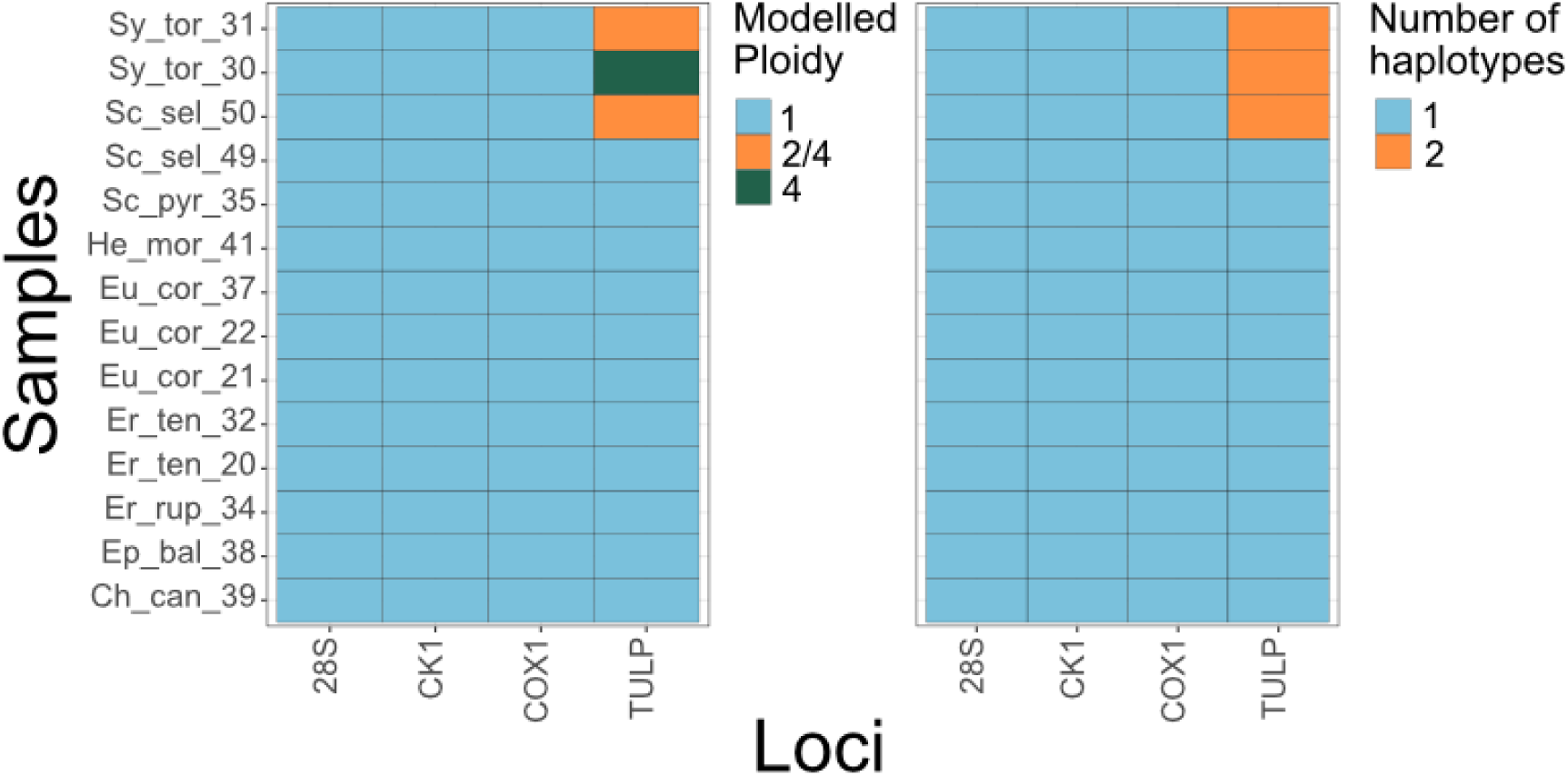
Heatmaps summarizing the consensus sequence reconstruction with amplicon_sorter. The left plot depicts the expected ploidy modelled for every sample and locus based on maximum likelihood. The right plot shows the total number of haplotypes, i.e. reconstructed consensus sequences that are used for downstream analyses.

The likelihood-based inference of putative ploidy-levels for samples and loci with more than one haplotype suggested that two heterozygous samples (Sy_tor_31 and Sc_sel_50) are most likely diploid and that reads that support the two haplotypes are present at a 50:50 ratio in the datasets for the TULP locus. Conversely, one sample (Sy_tor_30) appeared as tetraploid, since the ratio of the two haplotypes was close to 25:75. We, however, do not assume that this locus or sample is truly tetraploid, but rather that the TULP locus investigated here does not present a single-copy marker, but rather occurs in at least two paralogous gene copies as previously shown by Knippschild et al. (2005) and Ikeda et al. (2000).

Based on the reconstructed consensus sequences, we further generated locus-specific multiple alignments across all taxa. As shown in Figure S1 and according to our expectations, we found SNPs but no indel polymorphisms in the alignment of the mitochondrial COX1 locus. Conversely, we identified indel polymorphisms, putatively located in intronic regions, at several positions of the TULP locus. In particular, the outgroup bombyliid (He_Mor_41) was characterised by large insertions and several nucleotide changes. Consistent with these observations, we found different levels of sequence variability in pairwise comparisons of genetic differentiation at the four investigated loci, as shown in Figure 4, which indicates different evolutionary rates at the loci analyzed in this study. While indel polymorphisms are ignored in the phylogenetic inference based on maximum likelihood as implemented in IQ-tree2 (Minh et al., 2020), but we caution that misalignments around indels may lead to biased phylogenetic signals (Löytynoja & Goldman, 2008; Wong et al., 2008).

**Figure 4.**
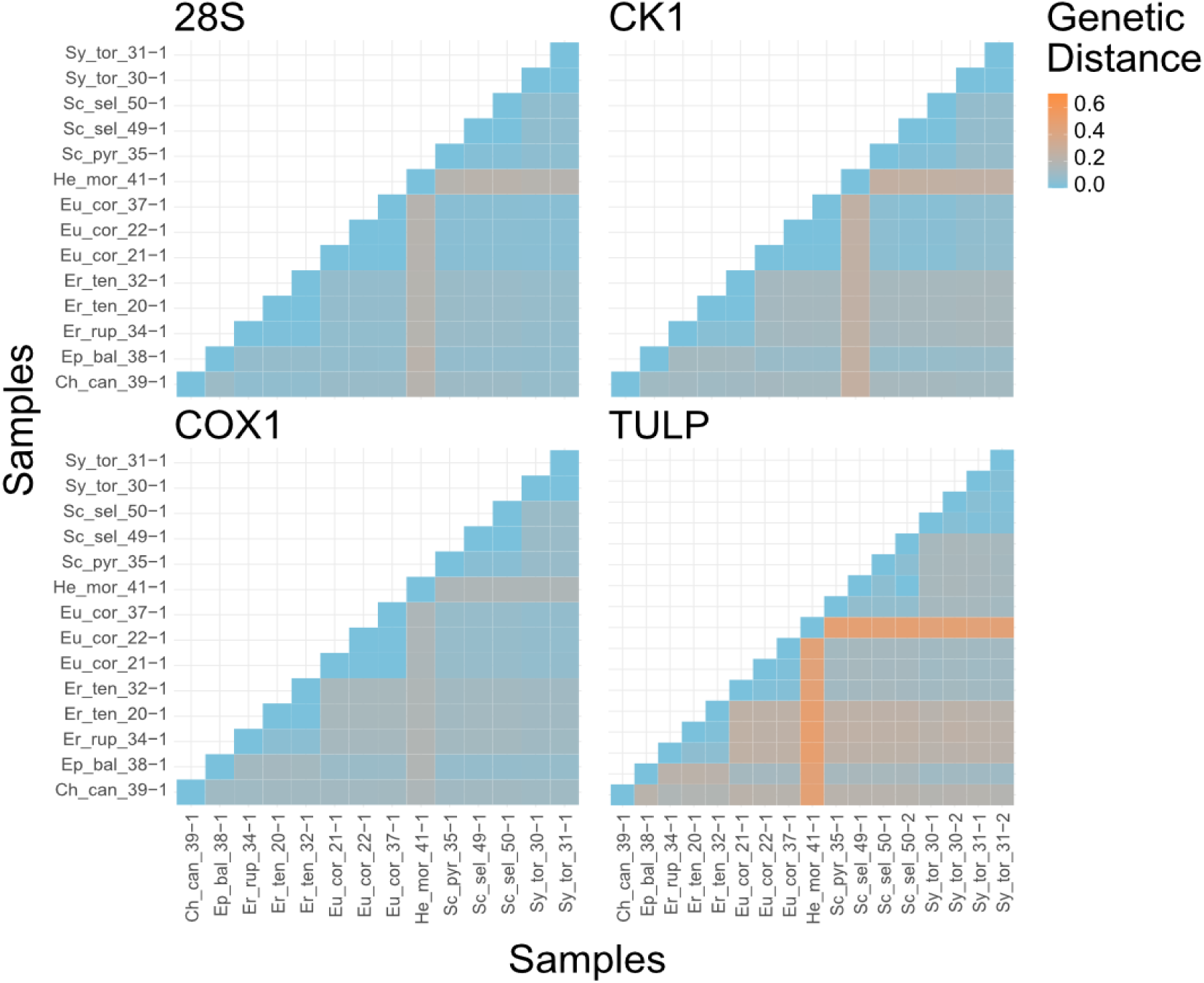
Heatmaps showing the amount of locus-specific genetic distances for all pairwise comparisons among all samples included in the dataset.

#### 3.1.2 Consistency of species identification based on morphology and DNA barcoding

We then used the reconstructed consensus sequences of each individual at the COX1 locus for comparison to the BOLD database. We inferred the putative species identity of each specimen based on sequence similarity to references in the database and found that all individuals matched in species identity to our previous identifications based on morphology (Table 3). All samples, except for one, were unambiguously assigned to the expected species in 10 out of the 10 top matches in the reference database. Sequence similarities were ranging from 92.05% (He_mor_41_1) to 100% for six specimens and only Ch_can_39_1 yielded four ambiguous species matches. However, the match with the highest sequence similarity (99.83%) correctly identified this specimen as *Cheilosia canicularis*, which is also the morphologically inferred species.

**Table 3.** Summary of the top 10 hits for species information obtained from BOLD for the consensus sequences for the COX1 locus.

| SAMPLE | Taxon1 (sim%) | Taxon2 (sim%) | Taxon3 (sim%) | Taxon4 (sim%) |
| --- | --- | --- | --- | --- |
| Er_ten_32_1 | *Eristalis tenax* (100.0%); count = 10 |  |  |  |
| Sc_sel_50_1 | *Scaeva selenitica* (99.97%); count = 10 |  |  |  |
| Eu_cor_21_1 | *Eupeodes corollae* (100.0%); count = 10 |  |  |  |
| He_mor_41_1 | *Hemipenthes morio* (92.05%); count = 10 |  |  |  |
| Ep_bal_38_1 | *Episyrphus balteatus* (100.0%); count = 10 |  |  |  |
| Er_rup_34_1 | *Eristalis rupium* (99.58%); count = 10 |  |  |  |
| Sy_tor_31_1 | *Syrphus torvus* (100.0%); count = 10 |  |  |  |
| Sy_tor_30_1 | *Syrphus torvus* (99.83%); count = 10 |  |  |  |
| Eu_cor_22_1 | *Eupeodes corollae* (99.9%); count = 10 |  |  |  |
| Eu_cor_37_1 | *Eupeodes corollae* (100.0%); count = 10 |  |  |  |
| Sc_pyr_35_1 | *Scaeva pyrastris* (99.64%); count = 10 |  |  |  |
| Ch_can_39_1 | *Cheilosia canicularis* (99.83%); count = 4 | *Cheilosia himantopus* (98.13%); count = 4 | *Cheilosia albipila* (94.65%); count = 1 | *Cheilosia impressa* (94.44%); count = 1 |
| Er_ten_20_1 | *Eristalis tenax* (100.0%); count = 10 |  |  |  |
| Sc_sel_49_1 | *Scaeva selenitica* (99.97%); count = 10 |  |  |  |

#### 3.1.3 Phylogenetic signal of different loci

Subsequently, we used the multiple alignment files for each locus separately, and for all loci combined, for phylogenetic inference using the maximum likelihood method as implemented in IQ-tree2 (v2.3.6, Minh et al., 2020). In addition, using the weighted ASTRAL method implemented in ASTER (v1.19.4.6, Zhang & Mirarab, 2022) we generated a joint super tree based on the unrooted trees of each locus. All trees in Figure 5 were rooted based on the bombyliid outgroup specimen He_mor_41_1 and showed inconsistent topologies for the individual loci. While individuals of the same genus were correctly clustered for all loci, the phylogenies based on the TULP and the COX1 loci could not be resolved and nested the cluster comprised of the subtribes Eristalinii (Er_ten_20_1, Er_ten_32_1 and Er_rup_34_1) and Rhingiini (Ch_can_39_1), without support for branching from bootstrapping (<75%), within the other specimens, which all belong to the Syrphini subtribe. Similarly, the phylogeny based on CK1 correctly placed the Eristalinii as a sister-group to the Syrphini but incorrectly placed the Rhingiini sample within the Syrphini. Only the 28S locus correctly resolved the expected phylogeny according to Moran et al. (2022). Moreover, both the ML tree based on the concatenated sequences of all loci and the ASTRAL tree (Figure 5 right) correctly resolved the phylogenetic relationships, which underlines that multilocus data is needed for unambiguous phylogenetic inference (e.g., Ermakov et al., 2015).

**Figure 5.**
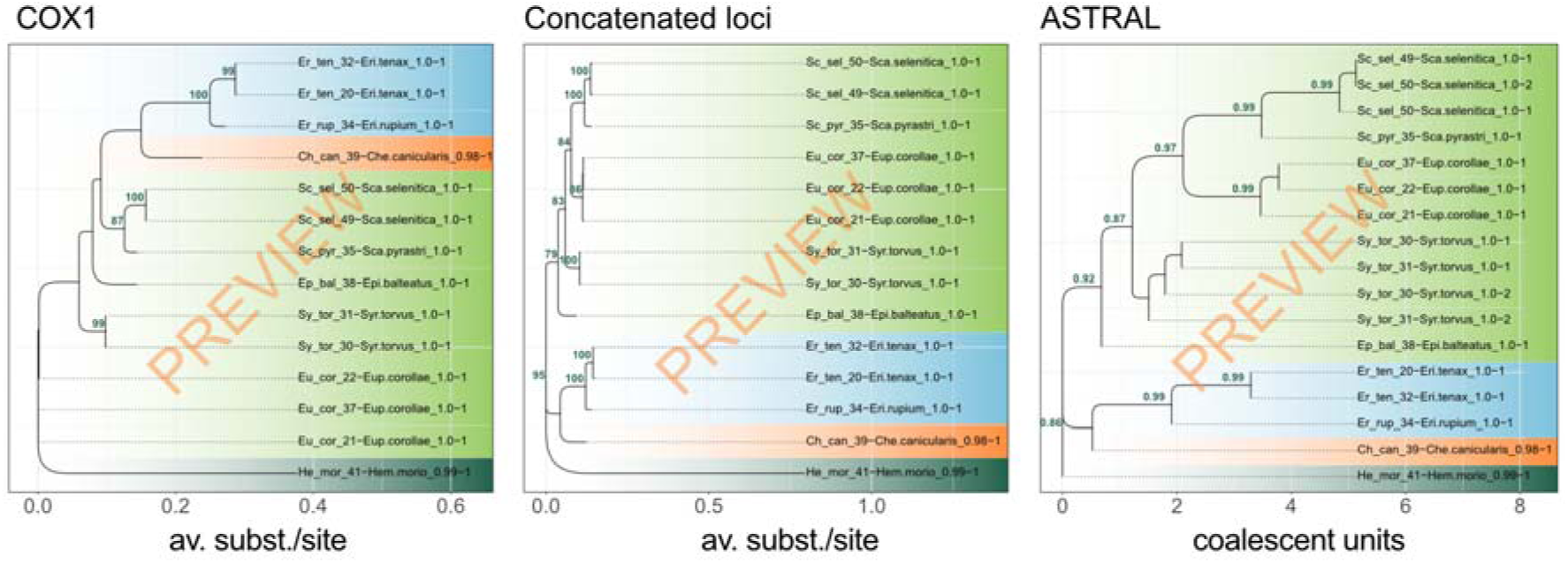
Phylogenetic trees based on maximum likelihood (1) using the sequence information of the COX1 locus only (left), (2) using the information of concatenated haplotypes across all loci (center) or based on a super tree generated with wASTRAL using the unrooted trees of each locus (right). All samples except for the bombyliid outgroup (dark green) belong to the subtribes Syrphini (light green), Eristalini (light blue) and Rhingini (orange).

#### 3.1.4 Species delimitation based on multiple individuals of different species

Finally, we used the MSA of the individual loci and the concatenated sequences for species delimitation with ASAP and found that the best models from the analyses based on several loci incorrectly collapsed either *Eristalis rupium* and *E. tenax* samples or *Scaeva pyrasti* and *S. selenitica* into single species (Figure 6). Moreover, consistent with the phylogenetic analyses presented above, which indicated that the TULP locus is characterized by only a few mutations, ASAP only distinguishes between the bombyliid outgroup and all syrphid samples in the best-fitting model. Conversely, we find that the combined dataset correctly distinguishes between species for all samples, which further underpins the diagnostic power of our multilocus approach.

**Figure 6.**
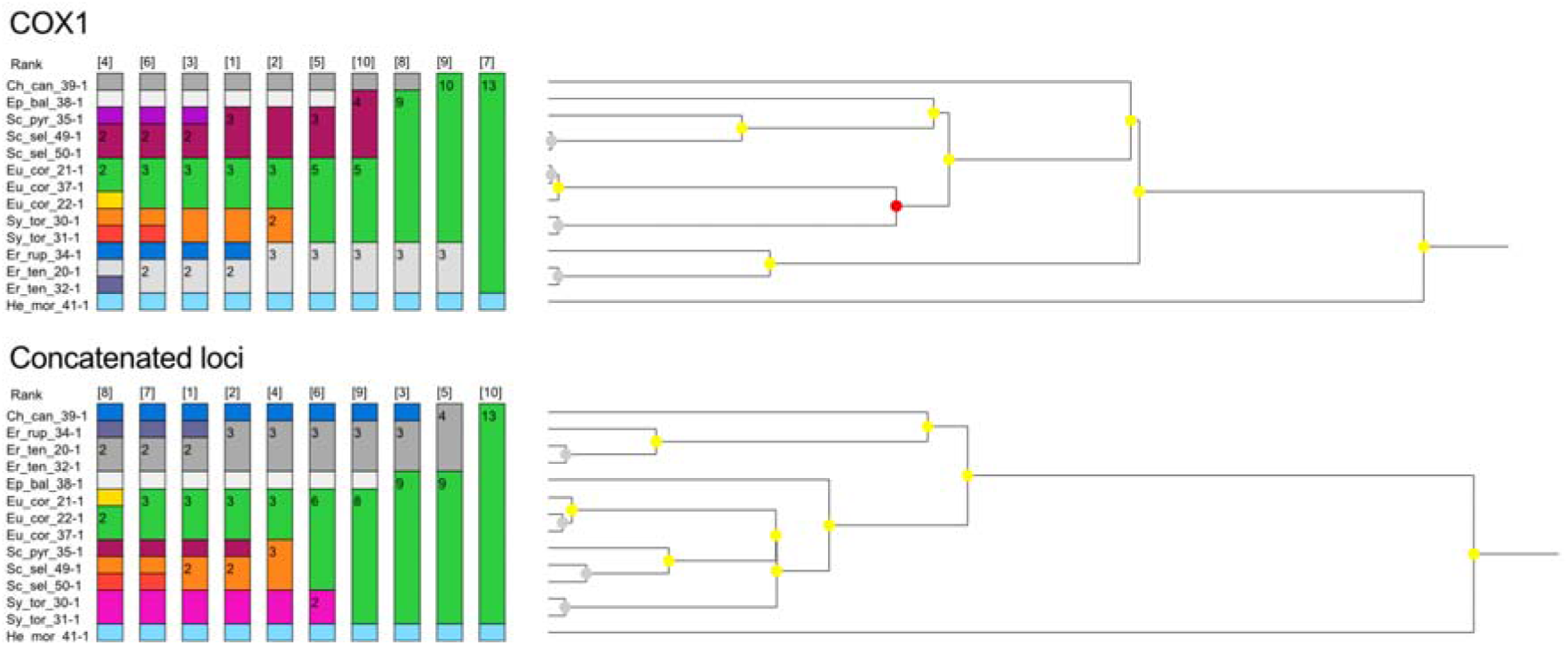
Grouping according to genetic distances based on hierarchical clustering as implemented in ASAP based on the COX1 sequence (top) or based on the concatenated sequences from all loci (bottom). The configuration with the best statistical support is indicated by the lowest number in each of the two analyses. The abbreviated specimen names correspond to the morphological species identification (see Table 1).

### 3.2 Benchmarking with real and simulated test datasets

The analyses based on ONT amplicon sequencing data from syrphid specimens, as presented above, allow us to test if the AmpliPiper results are consistent with previous data and biological expectations, such as species identity or phylogenetic signal. However, they cannot be used to assess the accuracy of the reconstructed consensus sequences since no reference sequencing data was available or the investigated specimens. To account for this, we employed a two-pronged approach.

First, we sequenced the haploid amplicons of the COX1 locus for all specimens in our dataset with Sanger sequencing and compared the sequences with the corresponding consensus sequences generated with AmpliPiper using different thresholds for demultiplexing and maximum read number for consensus reconstruction with amplicon_sorter. We found that our approach yielded identical sequences for all samples irrespective of the parameter combinations, which confirms that the consensus sequence reconstruction of our approach is very accurate and highly robust for haploid loci.

In a second step, we simulated Amplicon-Seq data based on reference sequences with differentiation levels ranging from 0% - 10% to further test the sensitivity and accuracy of our approach in reconstructing consensus sequences and in identifying different haplotypes in diploid samples. By combining simulated reads of two differentiated references at equal ratios we first tested if AmpliPiper is able to distinguish two haplotypes within a sample when using the default parameters for demultiplexing (*demultiplex.py, i.e.,--k-threshold 0.05 --min-read 1000*) while modifying the *--similar-consensus* (SC) parameter of amplicon_sorter, which influences at which similarity thresholds clusters of reads are collapsed during consensus sequence reconstruction. As shown in Figure 7 (top), we found a very strong influence of this parameter on the haplotype reconstruction of highly similar haplotypes. SC values of ≥ 97.5% and ≥ 98.5% correctly identified more than one haplotype at a sequence difference of 3% and 2%, respectively. However, SC values ≥ 98.5% occasionally (in one to two out of 20 technical replicates) incorrectly yielded three haplotypes irrespective of differentiation, which indicates that SC values between 97% and 98% provide the best results in terms of sensitivity. Larger SC values tend to be too sensitive and can result in spurious consensus sequences particularly at differentiation levels between 3% - 8% (see Figure 7).

**Figure 7.**
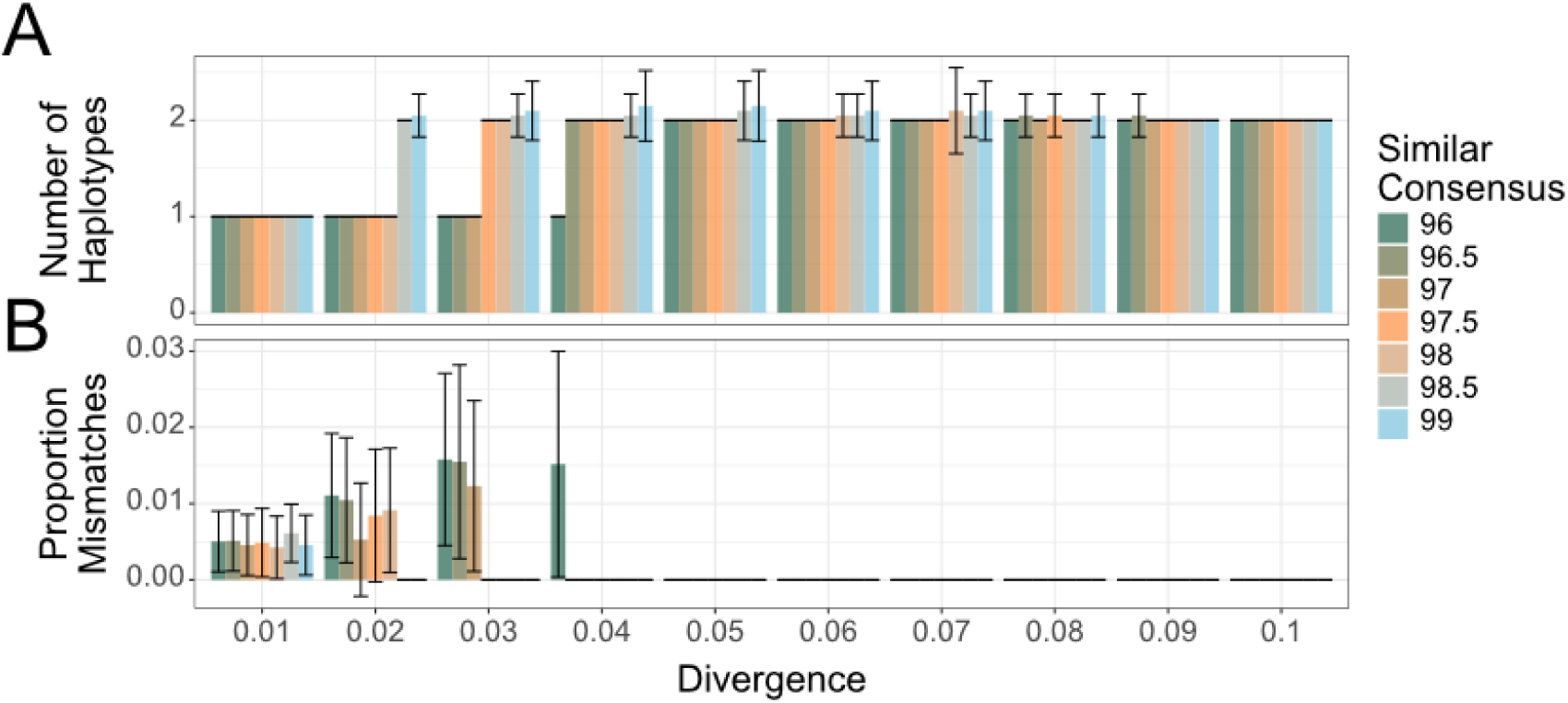
The barplots in panel A show the number of haplotypes, i.e. consensus sequences, that were reconstructed from equal proportions simulated reads that differed from 1% to 10% as shown on the x-axis. The barplots in panel B show the mean proportion of genetic differences of the reconstructed consensus sequence to the two differentiated original sequences. The colors indicate the value of the “similar consensus” parameter that was used for the analysis and which defines the threshold similarity among clusters of reads that are grouped into consensus sequences by amplicon_sorter. The error bars show standard deviations based on 20 technical replicates for each analysis.

To further assess the accuracy of our analyses, we compared the sequences of the reconstructed haplotypes to the original reference sequences. In cases where only one, rather than both haplotypes were successfully reconstructed, we found that the consensus sequences showed on average half the expected difference to the original reference (see Figure 7 bottom). Consistent with our expectations, these results indicate that these consensus sequences represent “collapsed” chimeric sequences intermediate in allelic composition between the two original haplotypes. Conversely, whenever both haplotypes were correctly reconstructed, we found that the consensus sequences were always identical to their respective reference sequences, which indicates that the consensus sequence reconstruction in our approach is very accurate even in diploid individuals.

Next, we tested if AmpliPiper is able to reconstruct haplotypes if their ratio deviates from 1:1, which may be the case, e.g., in tri- or tetraploid organisms. We therefore combined simulated reads from two references, where the frequency of the less frequent variant ranged from 10-50%. Consistent with results above, we found that haplotypes with differentiation levels <3% could not be distinguished irrespective of haplotype frequencies at an SC value of 97.5 (Figure S2). Conversely, AmpliPiper was able to correctly distinguish both haplotypes at differentiation levels ≥3% in most technical replicates even when the frequency of the minor haplotype was 10%, which indicates that the haplotype reconstruction is robust to unequal haplotype frequencies. Curiously, we found chimeric sequences in haplotypes with <3% difference only if both haplotypes occur at 1:1, but not if the ratios are unequal (with the exception of one case where a chimaera was reconstructed at a frequency ratio of 40:60 for two haplotypes with 1% difference).

Finally, we investigated the influence of read depth (RD) on the accuracy of the consensus sequence reconstruction. RD values of 10 and 20 were too low to yield any consensus (Figure S3). While we found that an RD of 50 was in most cases sufficient to reconstruct two consensus sequences, we observed that AmpliPiper failed to reconstruct both haplotypes in 1 out of 20 replicates at all differentiation levels. This suggests that RD > 50 are needed to yield robust results. Consistent with this conclusion, we found that RDs between 100 and 5000 had basically no influence on the accuracy for consensus reconstruction for haplotypes with differentiation levels ≥3%. In line with these observations, we found collapsed consensus sequences at differentiation levels of 1% and 2% irrespective of RD. Moreover, an RD of 50 resulted in chimeric sequences even at higher differentiation levels whenever only one haplotype was reconstructed.

### 3.3 Conclusion

In summary, the AmpliPiper analysis pipeline presented here integrates both cutting-edge, open-source third-party software and newly developed custom software to carry out a broad set of comprehensive analyses centered around multilocus DNA barcoding. Specifically, AmpliPiper uniquely employs demultiplexing of raw FASTQ files based on the expected fragment lengths and similarity of primer sequences to flanking regions of reads in pooled amplicon-seq data. These demultiplexed reads are then processed with amplicon_sorter to obtain sample- and locus-specific consensus sequences. Our benchmarking analyses based on comparisons of sample-specific consensus sequences for the COX1 locus generated with AmpliPiper to complementary Sanger sequences of the same samples revealed that all consensus and reference sequences were identical. This finding indicates that, in spite of the high error rate in ONT data, our combined approach of locus-specific demultiplexing of raw reads and consensus sequence reconstruction produces highly accurate consensus sequences. Moreover, we empirically assessed the sensitivity of our combined approach with simulated amplicon-seq data and showed that AmpliPiper is able to identify different haplotypes within a sample that differ by 3% mismatches. Our approach thus proves powerful enough to detect even within-sample variation and we are convinced that the sensitivity will further increase with future improvements of the ONT sequencing technology, which will further lower the sequencing error rate.

In contrast to other analysis software for amplicon-seq data which is usually tailored to specific analysis steps, AmpliPiper aims to provide a comprehensive yet user-friendly all-in-one solution that integrates several additional and follow-up analyses beyond consensus sequence reconstruction, which facilitate the interpretation and contextualization of the analyzed dataset. For example, AmpliPiper, is, to our knowledge, the only software which employs a maximum likelihood approach based on the frequency of the reads underlying the consensus sequences of different haplotypes to estimate the expected ploidy for a given locus and sample. Furthermore, AmpliPiper facilitates species identification for the widely used DNA barcode markers COX1, ITS and MATK_RBCL by utilizing the sequence information in the well-curated BOLD database or in GenBank to automatically obtain species information for reconstructed consensus sequences. Moreover, the subsequent phylogenetic and species delimitation analyses of all samples for each locus and across all loci combined allow a visual assessment and a critical biological interpretation of the data. Importantly, AmpliPiper provides detailed and well-structured output- and log-files which not only allow using the generated datasets for further analyses, but also to track errors and potential problems during the execution of the program.

However, DNA barcoding with multilocus approaches still has limitations that need to be considered when using AmpliPiper. For example, phasing multilocus haplotypes that are not physically linked is usually not possible. Likewise, AmpliPiper is not able to empirically phase multilocus haplotypes. Based on the outcome of amplicon_sorter, AmpliPiper labels different haplotypes for a given sample with consecutive numbers that are attached to the sample names. The order of the number is usually determined by the frequency of the haplotype in the pool of raw reads used by amplicon_sorter. Combined analyses across loci such as the super tree reconstruction with ASTER thus suffer from the problem that haplotypes with the same label from different loci are considered as phased, i.e. as being inherited from the same parent, and are jointly analyzed. This may be particularly problematic when investigating allopolyploid or hybrid organisms, which inherit whole genomes from different species (e.g., Gerchen et al., 2022). We therefore strongly recommend carefully assessing the genealogies in the ML trees of the individual loci, which may provide information about the hybrid nature of a given sample, prior to interpreting the ASTRAL tree and ML trees based on concatenated haplotypes. Moreover, our simulations have shown that the error rates in ONT sequencing data do not yet allow distinguishing haplotypes within a sample that differ in less than 3% mismatches. We recommend repeating the AmpliPiper analyses, at least on a subset of samples and loci, with different values of the similar consensus (--similar-consensus) parameters, similar to our analyses with simulated data, while retaining all reconstructed consensus sequences (--freqthreshold 0) to assess which parameter setting maximizes the sensitivity to identify any multiple haplotypes in nuclear markers. Given the possibility to comprehensively test different combinations of parameters and due to the simple visual depiction of the different analyses results, AmpliPiper is a useful tool to detect and visualize several possible confounding effects such as gene duplications, which may lead to unexpected ploidies, as found, for example, for the TULP locus in our dataset, but also DNA contaminants, which may lead to unexpected clustering of haplotypes with the contaminant samples. Thus, besides the main advantages of the AmpliPiper pipeline described above, supporting the evaluation of marker sequences for their suitability is a further strength of this approach. In summary, AmpliPiper aims to integrate results from multimarker approaches, where they are most needed: in cases where the reconstruction of taxonomic relationships is challenging due to a complex evolutionary history.

## Acknowledgments

This work has been funded by the TETTRIs project, which is funded by the Horizon Europe TETTRIs Grant Agreement 101081903. Astra Bertelli was funded by an Erasmus+ fellowship and Paula Schwahofer by the Österreichische Forschungsförderungsgesellschaft mbH (FFG, Project FO999903108: Syrphidae).

## 6 Data Accessibility and Benefit-Sharing Section

### 6.1 Data Accessibility

The (1) code of AmpliPiper including a detailed documentation, the (2) test datasets used to conduct the analyses presented in this manuscript (see testdata/) and the (3) analysis-pipeline for the simulations (see simulations/) can be obtained from https://github.com/nhmvienna/AmpliPiper.

### 6.2 Benefit-Sharing

Benefits generated: The analysis pipeline presented here is designed to facilitate integrative taxonomic and biodiversity research without advanced bioinformatics know-how. This will help to make the use of NGS more accessible for research in the fields of conservation biology and taxonomy. All specimens described in the publication have been collected with legal permits in Austria.

## 7 Authors contribution

Author contributions according to the CRediT’s contributor roles. AB: Conceptualization, Data curation, Formal analysis, Investigation, Methodology, Software, Validation, Visualization, Writing – original draft, Writing – review & editing; SS: Conceptualization, Data curation, Formal analysis, Investigation, Methodology, Software, Validation, Visualization, Writing – original draft, Writing – review & editing; SK: Investigation, Methodology, Validation, Writing – review & editing; PS: Formal analysis, Investigation, Writing – review & editing; EH: Conceptualization, Funding acquisition, Resources, Writing – review & editing; NS: Conceptualization, Funding acquisition, Resources, Writing – review & editing; LK: Conceptualization, Funding acquisition, Resources, Supervision, Writing – review & editing; LK: Conceptualization, Funding acquisition, Resources, Supervision, Writing – review & editing; MK: Conceptualization, Data curation, Formal analysis, Funding acquisition, Investigation, Methodology, Project administration, Software, Supervision, Validation, Visualization, Writing – original draft, Writing – review & editing

## Supplementary Figures

**Figure S1.**
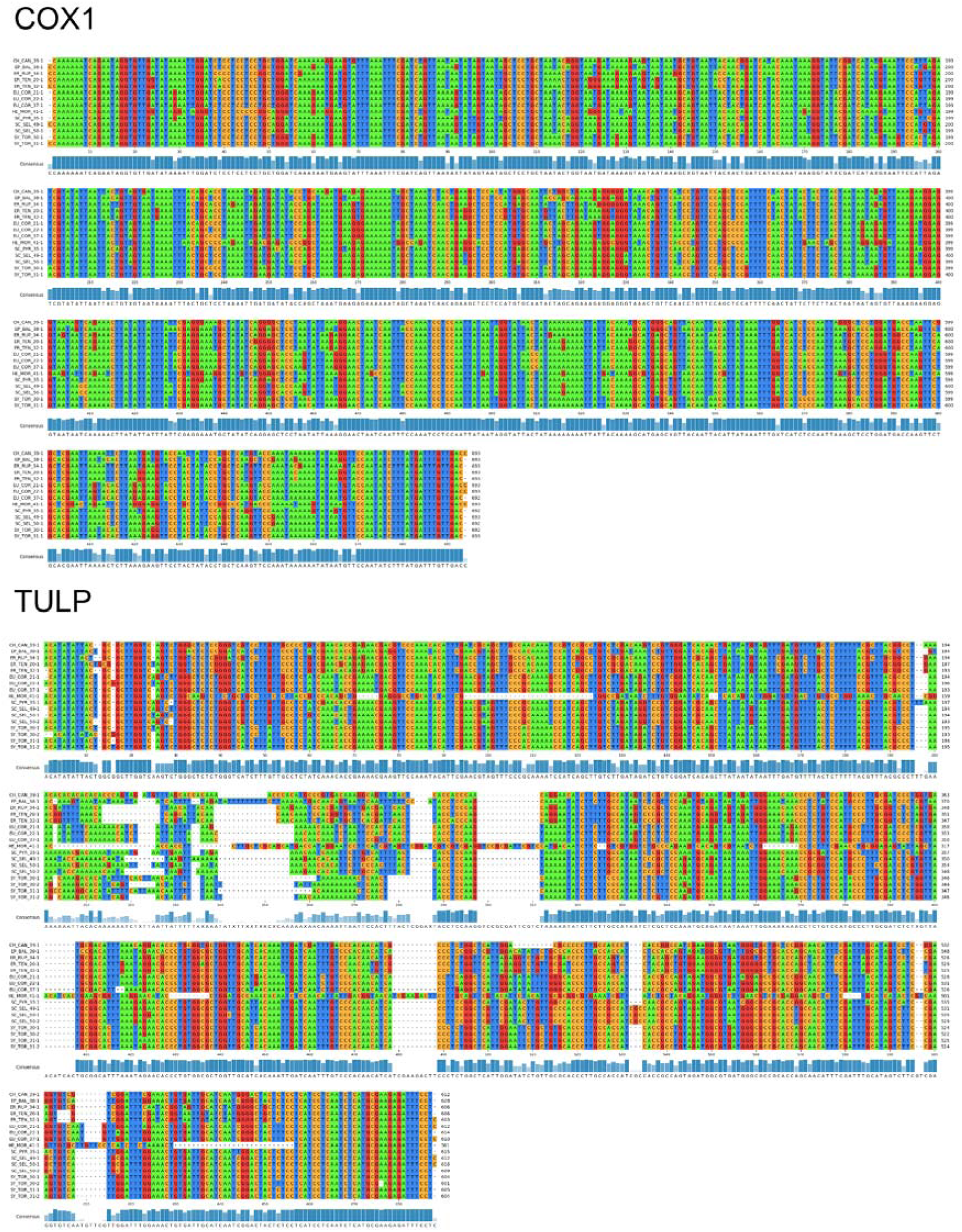
Multiple alignment of consensus sequences of the COX1 (top) and the TULP (bottom) sequences.

**Figure S2.**
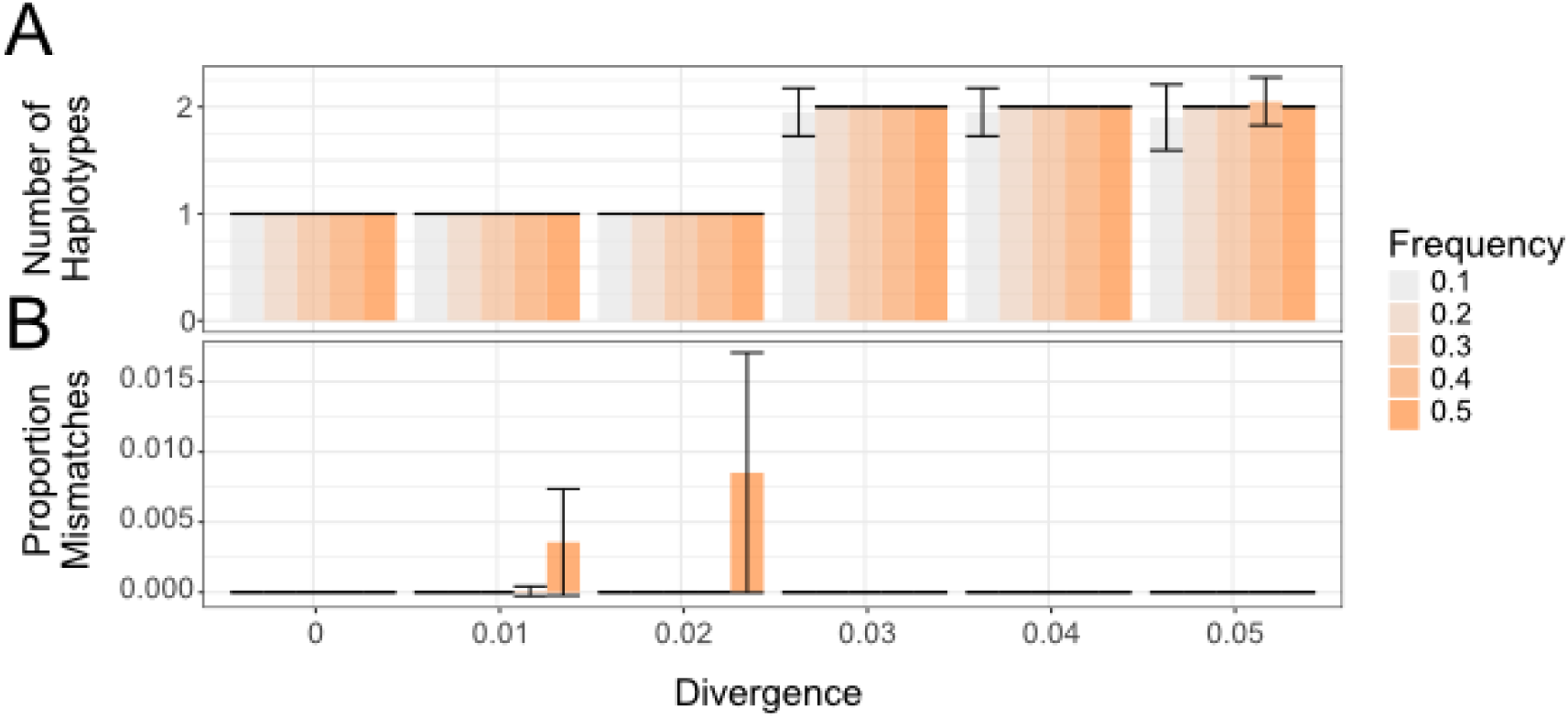
The barplots in panel A show the number of haplotypes, i.e. consensus sequences that were reconstructed from different proportions of simulated reads that differed from 0% to 5% as shown on the x-axis. The barplots in panel B show the mean proportion of genetic differences of the reconstructed consensus sequence to the two simulated reads in the pools. The colors indicate the proportion of the reads with the lower frequency in the pool. The error bars show standard deviations based on 20 technical replicates for each analysis.

**Figure S3.**
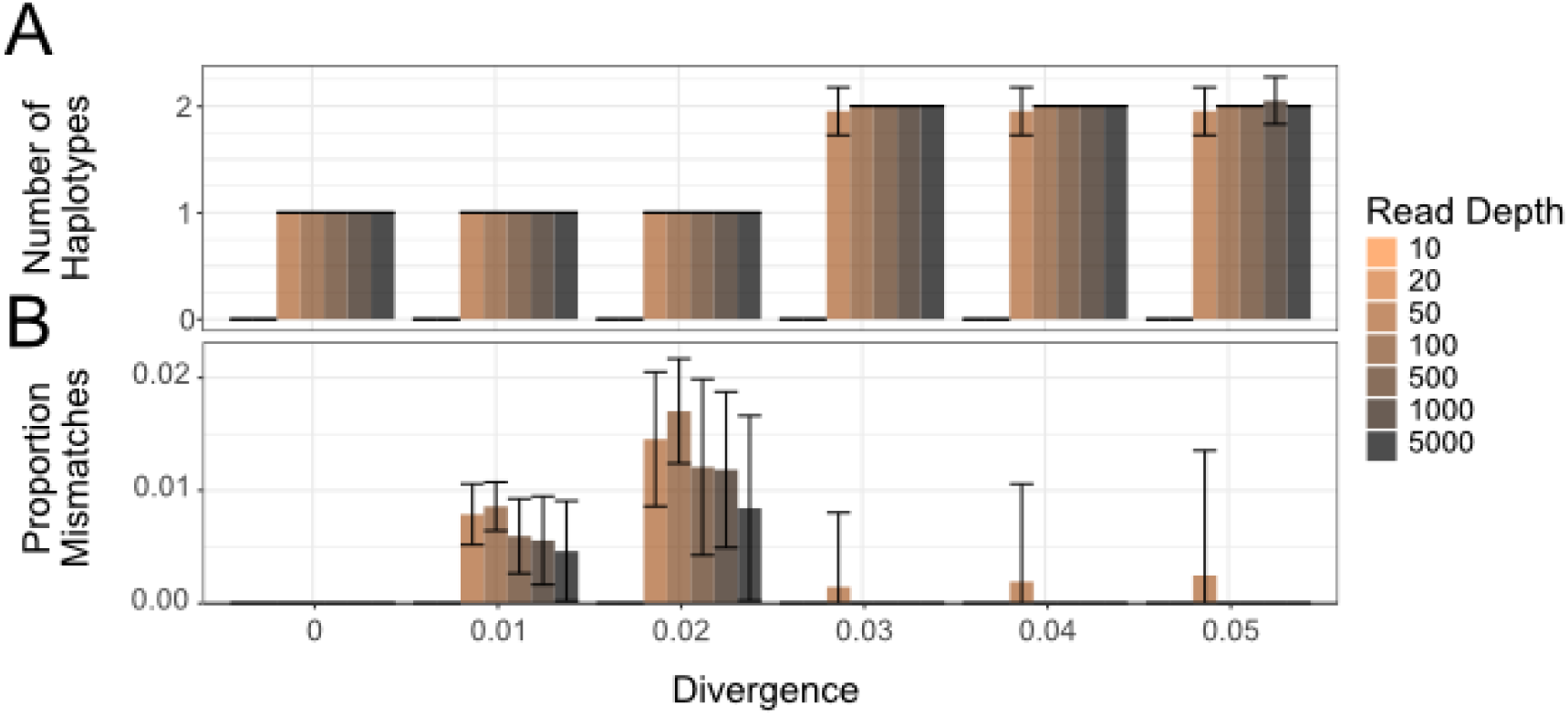
The barplots in panel A show the number of haplotypes, i.e. consensus sequences, that were reconstructed from equal proportions of simulated reads that differed from 0% to 5% as shown on the x-axis. The barplots in panel B show the mean proportion of genetic differences of the reconstructed consensus sequence to the two simulated reads in the pools. The colors indicate the total number of pooled reads provided to amplicon_sorter. The error bars show standard deviations based on 20 technical replicates for each analysis.

